# Evolution of Phenotypic Plasticity under Host-Parasite Interactions

**DOI:** 10.1101/2021.01.20.427416

**Authors:** Naoto Nishiura, Kunihiko Kaneko

## Abstract

Robustness and plasticity are essential features that allow biological systems to cope with complex and variable environments. Through the evolution of a given environment, the former, the insensitivity of phenotypes, is expected to increase, whereas the latter, the changeability of phenotypes, tends to diminish. However, in nature, plasticity is preserved to a certain degree. One possible cause for this is environmental variation, with one of the most important “environmental ” factors being inter-species interactions. As a first step toward investigating phenotypic plasticity in response to an ecological interaction, we present the study of a simple two-species system consisting of hosts and parasites. Hosts are expected to evolve to achieve a phenotype that optimizes fitness and increases the robustness of the corresponding phenotype by reducing phenotypic fluctuations. Conversely, plasticity evolves in order to avoid certain phenotypes being attacked by parasites. By simulating evolution using the host gene-expression dynamics model, we analyze the evolution of genotype-phenotype mapping. If the interaction is weak, the fittest phenotype of the host evolves to reduce phenotypic variances. In contrast, if a sufficient degree of interaction occurs, the phenotypic variances of hosts increase to escape parasite attacks. For the latter case, we found two strategies: if the noise in the stochastic gene expression is below a certain threshold, the phenotypic variance increases via genetic diversification, whereas above the threshold, it is increased due to noise-induced phenotypic plasticity. We examine how the increase in the phenotypic variances due to parasite interactions influences the growth rate of a single host, and observed a trade-off between the two. Our results help elucidate the roles played by noise and genetic mutations in the evolution of phenotypic plasticity and robustness in response to host-parasite interactions.

**Author summary:** Plasticity and phenotypic variability induced by internal or external perturbations are common features of biological systems. However, certain environmental conditions initiate evolution to increase fitness and, in such cases, phenotypic variability is not advantageous, as has been demonstrated by previous laboratory and computer experiments. As a possible origin for such plasticity, we investigated the role of host-parasite interactions, such as those between bacteria and phages. Different parasite types attack hosts of certain phenotypes. Through numerical simulations of the evolution of host genotype-phenotype mapping, we found that, if the interaction is sufficiently strong, hosts increase phenotypic plasticity by increasing phenotypic fluctuations. Depending on the degree of noise in gene expression dynamics, there are two distinct strategies for increasing the phenotypic variances: via stochasticity in gene expression or via genetic variances. The former strategy, which can work over a faster time scale, leads to a decline in fitness, whereas the latter reduces the robustness of the fitted state. Our results provide insights into how phenotypic variances are preserved and how hosts can escape being attacked by parasites whose genes mutate to adapt to changes in parasites. These two host strategies, which depend on internal and external conditions, can be verified experimentally, for example, via the transcriptome analysis of microorganisms.

## Introduction

Robustness and plasticity are two important properties of biological systems. To maintain function and high fitness in response to internal noise, environmental variation, and genetic changes, the fitted state must be robust, whereas the phenotype needs to possess plasticity in order to adapt to environmental variation. Indeed, the evolution of robustness and plasticity has been investigated extensively both theoretically and experimentally [1–5].

In particular, the relationship between phenotypic fluctuations and robustness or plasticity has recently attracted much attention [6]. As a result of evolution under a given environment, phenotypic fluctuations generally decrease. With this decrease, the system deviates less from the fitted state, thereby increasing robustness [7–9]. However, decreased phenotypic fluctuation is also accompanied by decreased adaptability to environmental variation, is discussed by generalising the fluctuation-response relationship in statistical physics [6, 10, 11]. Here, phenotypes are subject to the dynamic processes of several gene-dependent variables (for example, the expressions of proteins). Thus, phenotypes vary according to genetic changes, providing the phenotypic fluctuations [12–14]. On the other hand, as the dynamic processes that shape phenotypes are influenced by external and internal noise, phenotypes can vary even for isogenic individuals [15–17], representing another source of phenotypic fluctuations [18, 19]. Both these fluctuation sources depend on genotype-phenotype mapping [20–22]. The evolution of genotype-phenotype mapping is essential to understand the evolution of robustness and plasticity, and has been explored extensively both in numerical [4, 6] and laboratory experiments [23, 24].

Indeed, experiments and simulations of adaptive evolution under fixed conditions have shown that fluctuations decrease over the course of evolution [25], and the evolvability, that is, the rate of increase in the fitness or the phenotypic change per generation, declines. Accordingly, as evolution progresses, the robustness of the phenotype increases, while the plasticity decreases.

However, in nature, phenotypic fluctuations and evolutionary potentiality persist. Even after evolution, the phenotype in question is not necessarily concentrated on its optimal value, but its variance often remains rather large. For instance, even for isogenic cells (clones), fluctuations in the concentration of each protein remains sufficiently high to preserve potentially of evolution. This leads to ask the following questions to be addressed: Why are such fluctuations not reduced, which, in principle, would be possible by evolving appropriately negative feedback processes for stabilization? How are phenotypic fluctuations or plasticity sustained?

One possible cause for the preservation of plasticity or fluctuation is environmental fluctuation [26–28]. In natural evolution, fitness and environment are not fixed, but variable. If environmental fluctuation is reduced excessively, the plasticity to adapt to new conditions imposed by environmental changes would be lost. The plasticity of a biological system dictates its capacity to cope with environmental changes that alter fitness conditions.

For every species, interactions with other species are one of the most important “environmental” factors that affect its existence. Even if a certain “external” environmental condition itself is fixed, the types and populations of other species may change owing to species-species interactions [29]. For instance, in the interaction between hosts (prey) and parasites (predators), the former may change their phenotype to escape attack by the latter, which, in turn, will change the phenotype of the latter to continue attacking the former [30, 31]. Thus, each species may retain evolutionary plasticity to cope with dynamic changes in other species. Furthermore, the phenotypic plasticity in one species may influence other species, resulting in a dynamic change in inter-species interactions. If sufficient species-species interaction occurs, the dynamic variation of phenotypes may be mutually sustained across multiple species.

As a first step to investigate the phenotypic plasticity originating from ecological interactions, we present a study of a simple two-species system consisting of hosts and parasites. Within this system, distinct parasite types exist that can attack hosts possessing a specific phenotype, whereas the host phenotypes are determined by genotype-phenotype mapping designed to incorporate possible fluctuations. Hosts are expected to evolve to achieve a phenotype that optimizes fitness and increases the robustness of the successful phenotype by reducing phenotypic fluctuations. In addition, they are also expected to evolve plasticity, that is, phenotypic adaptability to cope with parasite attacks. We use a host gene-expression dynamics model to study the evolution of genotype-phenotype mapping and the resultant phenotypic fluctuations, subsequently exploring the evolution of phenotypic robustness and plasticity in response to interactions with parasites. Specifically, we address the following questions: first, how are the robustness and plasticity of phenotypes, which tend to exhibit opposite trends, maintained through evolution due to the interaction between host and parasite species? Second, are phenotypic fluctuations sustained to cope with parasite attacks? Here, as mentioned, there are two sources of phenotypic fluctuations: genetic variation and stochasticity in genotype-phenotype mapping (that is, noise in the dynamics that shape the phenotype). Then, we address a third question: which of these sources of phenotypic fluctuation is dominant in sustaining the level of phenotypic fluctuation? Finally, we discuss the dependence of the evolution of phenotypic fluctuation and plasticity on the strength of the host-parasite interaction as well as the sensitivity of phenotype dynamics to noise.

## Models

### Gene regulatory networks

To study the evolution of genotype-phenotype mapping, we employed a simple model for gene expression dynamics based on a sigmoidal function. This model comprises a network consisting of genes that mutually activate or inhibit each expression. In this model, there are *M* genes whose gene expression level, 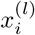 (*i* = 1, 2, …, *M*), corresponding to the host type *l* at time *t* is described by

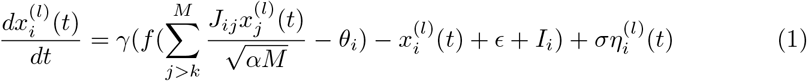

where *J*_*ij*_ is the matrix representing the influence of gene *j* on the expression of gene *i*. When *j* activates (represses) the expression *i*, it takes the form *J*_*ij*_ = 1(−1), where *J*_*ij*_ = 0 if *j* does not influence *i*. The sigmoidal function, *f*(*z*), is given by

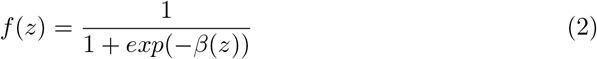

for which *β* = 25, meaning that *f*(*z*) closely resembles a step function. Furthermore, *θ*_*i*_ represents the threshold of the input required for the expression of gene *i*. Therefore, whether the gene is expressed (*f*(*z*) ≈ 1) or not (*f*(*z*) ≈ 0) depends on whether the sum of inputs from the expression of other genes is larger than *θ*_*i*_. Moreover, there are *k* “ output genes “ *i* = 1, 2, …, *k*, which determine the fitness as defined below. In Eq (1), the value *θ*_*i*_ is distributed randomly between fixed limits of 0.05 and 0.3. The term representing the interaction with other genes scales with 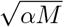, where α is the path density and the fraction of non-zero values of *J*_*ij*_. We adopted this scaling so that *x*_*i*_ is comparable with the order of *θ*_*i*_ regardless of *M*. To eliminate the potential for the output genes to influence others, the summation is taken only for *j* > *k*. The term *η*^(*l*)^(*t*) denotes a Gaussian white noise originating from molecular fluctuations in chemical reactions and satisfying< *η*_*i*_(*t*)*η*_*j*_(*t*) >= *δ*_*i,j*_*δ*(*t* − *t′*), which represents the stochasticity. The value of *σ* represents the strength of this noise. Initially, all the gene expression levels are set smaller than the threshold *θ*_*i*_. Furthermore, *ϵ* denotes the spontaneous expression level, which is smaller than *θ*_*i*_, and dictates that *x*_*i*_ must receive external or internal inputs from other *x*_*j*_ values in order to advance beyond *θ*_*i*_. *I*_*j*_ is the external input for” input genes, “ with *I*_*j*_ = 1 for *j* = *M* − *l*_*inp*_ + 1, *M* − *l*_*inp*_ + 2, …, *M*, and *I*_*j*_ = 0 for *j* ≤ *M* − *l*_*inp*_. Initially, we choose *J*_*ij*_ at random to obtain *J*_*ij*_ = 1(−1) with a probability of 0.15, in order that α = 0.3. Additionally, we set *N* = 300, *M* = 64, *l*_*inp*_ = 8, and *l*_*out*_ = 8.

### Fitness and reproduction

The fitness of the host alone i.e., in the absence of parasites is determined by the fraction of output genes that are expressed, as ds

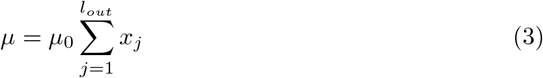

where *x*_*j*_ is the expression of the output genes and µ_0_ denotes the maximum fitness. Therefore, the state with all output genes expressed (*x*_*j*_ = 1, *j* = 1, 2, …, *l*_*out*_) is an optimal phenotype for achieving maximal growth.

Next, we introduce parasite attacks into the model. There are 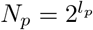 parasites, coded as 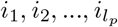 with *i*_*j*_ = 0 or 1, such that 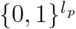. For instance, for *l*_*p*_ = 3, there are 2^3^ = 8 species coded by 000, 001, …, 111. Each parasite attacks hosts that possess the corresponding target gene expression, 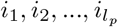, that is, each parasite attacks the host whose gene expression pattern, *x*_*j*_ (1 ≤ *j* ≤ *l*_*p*_), matches with 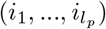. Below, we term the *l*_*p*_ genes as” target genes. “Here, the target condition is given by *s*(*x*_*i*_) = 1 for *x*_*i*_ > 0.5 and *s*(*x*_*i*_) = 0 otherwise (as the threshold function is close to a step function, *x*_*i*_ takes ∼ 0 or ∼ 1 any way). For instance, a host whose expression pattern (*s*(*x*_1_), *s*(*x*_2_), *s*(*x*_3_)) ≈ (0, 1, 0) is attacked by the parasite (*i*_1_, *i*_2_, *i*_3_) = (0, 1, 0). The growth rate, µ_*i*_, of each host decreases owing to the impact of the parasite *i* as follows:

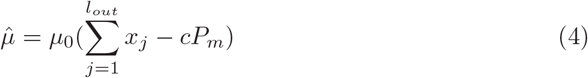

where *P*_*m*_ is the population density of the m-th parasite 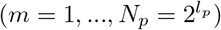 and c is the strength coefficient of the attack by the parasite. The volume of the host cells grows according to

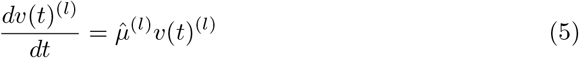

When the volume exceeds a threshold value of 2 (*v*(0) set to 1), the host cell is assumed to divide into two parts. If cell division leads to the cell number exceeds the upper limit *N* by the cell division, the surplus cells are eliminated at random, in order that the upper limit of the total cell number is *N*. Here, after division, the initial expression of *x*_*i*_ is reset to take a random value smaller than the threshold *θ*_*i*_. Mutation is factored into the division process and added to the network *J*_*ij*_ with a low mutation rate. An (*i, j*) path in the network matrix is selected at random and its values changed to one of the other to preserve the total path number. For example, if *J*_*ij*_ = 1(−1) and *J*_*kj*_ = 0, then they are changed to *J*_*kj*_ = 1(−1) and *J*_*kj*_ = 0. We prohibited direct connections between the “input” and “output” genes.

### Parasite population dynamics

The population of each parasite *P*_*i*_ increases in proportion to the host population with the corresponding phenotype, where the host density, *H*_*i*_, to be attacked by the parasite strain, *i*, is determined by the host population that satisfies *s*(*x*_*j*_) = *i*_*j*_. In addition, there is a mutation from other parasite types 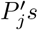. Because parasite types are represented by a binary string of length *l*_*p*_, they are represented in *l*_*p*_-dimensional hypercube space. Each distinct genotype 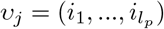 is represented by a vertex on the cube. Mutations in the parasite species occur by flipping each gene *i*_*l*_ = 0 ↔ *i*_*l*_ = 1 at a rate of *D*_*p*_. This is represented by diffusion in the hypercube (e.g., 010 ↔ 000) determined by a diffusion constant *D*_*p*_. Therefore, the population dynamics is expressed as

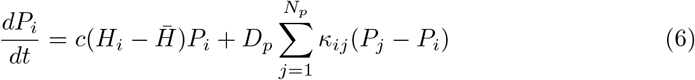

where *κ*_*ij*_ takes unity only if the binary string of *j* can change to that of *i* via a single mutation, otherwise it equals zero. To avoid the divergence of parasite populations, we assume competition within all the parasite species, such that the total number of populations is assumed to be fixed. Accordingly, we normalize the density *P*_*i*_ such that 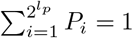. Thus, the reduction term 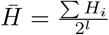 is introduced ensuring that 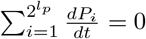 and 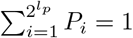 remain satisfied.

### Phenotypic variances due to noise and mutation

The model includes a noise component that allows the fitness to fluctuate during each run, which leads to the distribution in the expression of *x*_*i*_, even among individuals sharing the same gene regulation network. We define the phenotypic variance *V*_*p*_(*i*) as the variance of the phenotype *x*_*i*_ over the host population with different genotypes. In contrast, we compute the variance 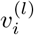 of the gene expression of 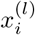 for each host type *l*. Because the model includes this noise term, the variance is finite, and the expression level *x*_*i*_ for the same host genotype (network) varies between individuals. Here, the variation of *x*_*i*_ to give *V*_*p*_(*i*) arises both from noise and from network mutations. To distinguish between these sources, we define two phenotypic variances: *V*_*ip*_(*i*) and *V*_*g*_(*i*) [9, 10]. The variance 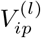 is defined as the variance of phenotypic fluctuations in the isogenic population, that is, the phenotypic variance of *x*_*i*_ within the clonal population of the host type *l*. In contrast, 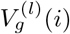 is defined as the phenotypic variance due to the genetic distribution, as computed by the variance of 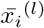 within the heterogenic population and adding a mutation to 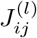, where 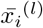 is the average over the clonal distribution of host *l*. Details describing the calculation of the variances *V*_*ip*_ and *V*_*g*_ are provided in the *Methods* section. Here, we are especially interested in the variance of the expression of output genes. We term the average variances of the output genes (*i* = 1, 2, …, 8) as *V*_*ip*_, *V*_*g*_, and *V*_*p*_, thereby omitting the *l* notation. Additionally, we define the average variances of the target genes corresponding to parasite attacks as 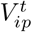 and 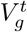.

## Results

### Parasite interaction accelerates host diversity

Fig 1 shows the population dynamics and fitness values of the host in three parasite environments (*c* = 0, 1, and 4). In the absence of parasite interaction, i.e., at *c* = 0, gene regulatory networks (GRNs) that generate a phenotype expressing all target genes, such as type“111” (black line), dominate the host population (see Fig 1(i)). Genotypes that lead to the fittest phenotype that expresses all the output genes dominate in the population. Then, if sufficient interactions with the parasites occur, the host population evolves into multiple groups with different phenotypes (Fig 1(ii), (iii)). This is explained by the selection pressure to avoid parasite attacks. When the host population is concentrated on the fittest type “11111111,” which includes the target gene expressions “111,” the parasite population is also concentrated on the corresponding type matching the target “111,” resulting in the population of the fittest host types being suppressed by the interaction with the parasite. Consequently, other host types with less fitted target patterns have more chances to survive. In short, parasites act as a negative frequency-dependent selection pressure, and the effective fitness (Eq (4)) varies over time owing to the change in the parasite population distribution of each type. As a result, multiple phenotypes and genotypes can coexist within a host population.

**Fig 1.**
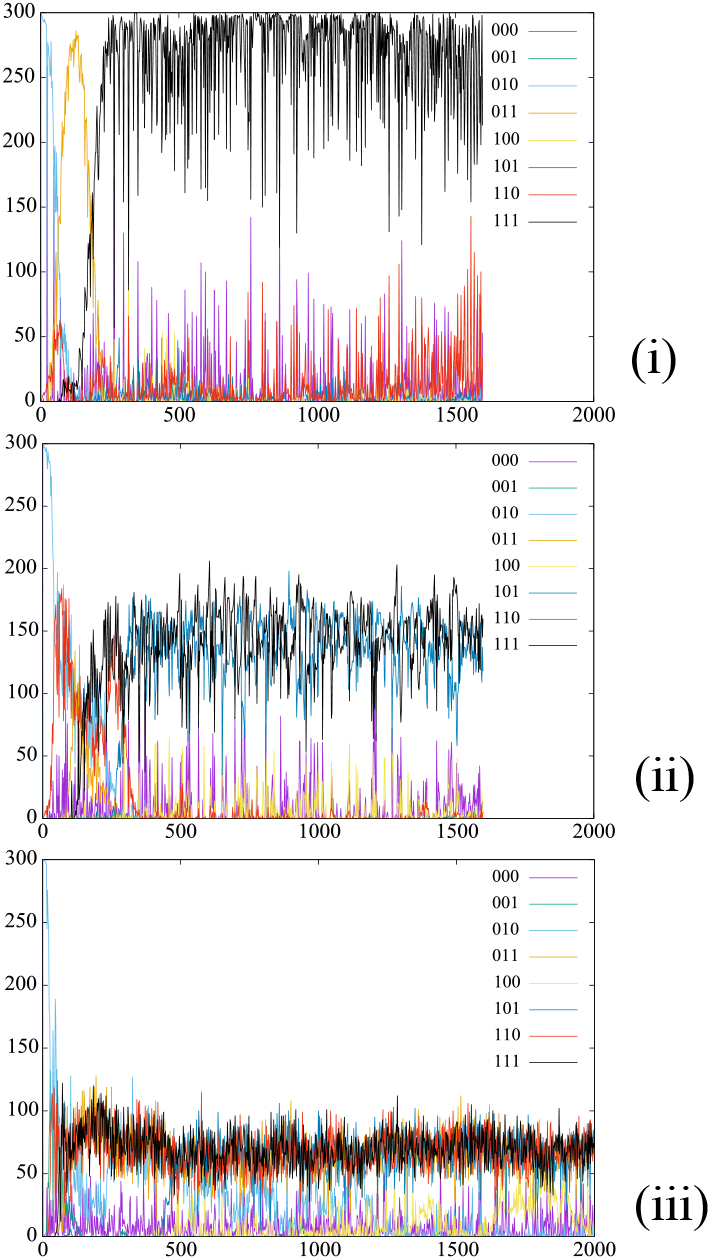
Dynamics of host phenotypes under interaction with parasites. The populations of hosts *H*_*i*_ corresponding to the parasite type 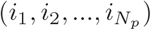 are plotted against the generation time. The number of parasite types is *N*_*p*_ = 2^3^ = 8, which corresponds to the on/off expression of three target genes of the host. The population of each host type *i* for *H*_*i*_ representing *s*(*x*_1_), *s*(*x*_2_), *s*(*x*_3_) =“ 000”, “ 001”,…, and” 111” is indicated by different colors. The interaction strength is *c* = 0 (i), 1 (ii), and 4 (iii). In the absence of parasites (*c* = 0), the population is dominated by the “111” type in which all output gene expression levels are turned on, whereas other phenotypic types with low growth rates (“ 110,” “101,” and” 011”) also coexist at *c* = 1, 4. In addition, *N* = 300, M = 64, *l*_*inp*_ = 8, *l*_*out*_ = 8, and *l*_*p*_ = 3 (see Table 1).

To determine the extent to which host-parasite interactions enhance phenotypic diversity, we computed the phenotypic variance, *V*_*p*_, that is, the variance of output gene expressions over the host population with distributed genotypes (i.e., network *J*_*ij*_) and plotted it as a function of the interaction strength *c*, as shown in Fig 2. These variances were computed with respect to the time-dependent average of the output gene expression levels, *x*_*i*_. As the interaction strength *c* is larger than the threshold *c*_*t*_ = 1, *V*_*p*_ exhibits a sharp rise. At *c* ∼ *c*_*t*_, a transition is observed, beyond which the phenotypic diversity increases. This diversification means that multiple phenotypes coexist within a group, as shown in Figs. 1(ii) and (iii). In contrast, below the transition point, *c*_*t*_, the variance decreases and the host population is concentrated on the fittest phenotype, as shown in Fig 1(i).

**Table 1.** Model parameters and variable definitions

| Symbol | Description | value |
| --- | --- | --- |
| $M$ | total number of host genes | 64 |
| $N$ | total number of host cells | 300 |
| $l_{inp}$ | number of input genes | 8 |
| $l_{out}$ | number of output genes | 8 |
| $I_j$ | input strength | -5 |
| $\gamma$ | time constant | 0.1 |
| $\alpha$ | average path density | 0.1 |
| $\epsilon$ | spontaneous expression level | 0.01 |
| $\beta$ | | 25 |
| $\mu_0$ | maximum growth rate | 0.0002 |
| $l_p$ | total number of parasite genes | 3 |
| $N_p$ | number of parasite types | 8 |
| $T_p$ | evolutionary speed | 100 |

**Fig 2.**
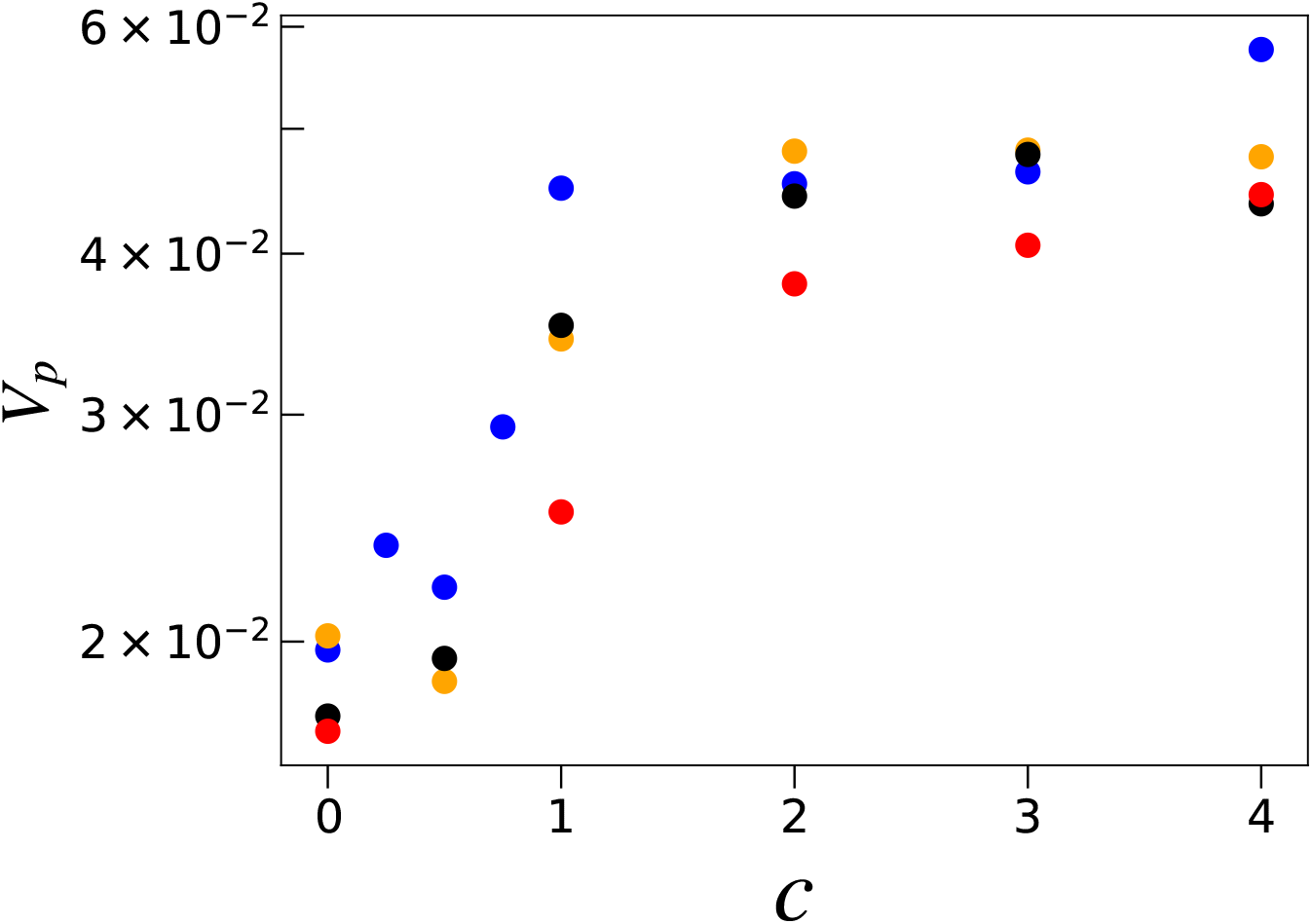
Total phenotypic variance *V*_*p*_ of the output genes against the parasite interaction strength. The variance *V*_*p*_ of the output genes plotted against the parasite interaction strength *c. V*_*p*_ is computed from the expression levels of the output genes over the host population (*N* = 300 individuals) and plotted as the average *V*_*p*_ over 2500–3000 generations. Each color represents a different noise strength: σ = 0.03 (blue), 0.02 (orange), 0.01 (black), and 0.0 (red).

This transition point for phenotypic diversification is determined from the fitness function, as defined in Eq (4). The growth rate is maximized when the expression level of all the output genes is close to *x*_*j*_ = 1. In the present model, the fitness decreases by one unit when one of the output genes is switched off. Furthermore, the maximum increase in the growth rate required to escape the parasite attack by changing phenotype is given by *c*. Thus, at *c* ≈ 1, the benefit of avoiding the parasite attack by switching off one of the target genes from the fittest type exceeds the associated reduction in the growth rate. Therefore, the transition point *c*_*t*_ is estimated by *c* ≈ 1. Indeed, Fig 2 shows that, for *c* > *c*_*t*_, the variance increases. When the interaction is strong (e.g., *c* = 4), the number of host phenotypes approaches the maximum number, 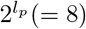, corresponding to all possible parasite species (see Fig 1(iii)). Note that the gene expression dynamics involve noise; therefore, hosts of the same genotype can exhibit different expression patterns among individuals of the same genotype. Accordingly, phenotypic variation has two different origins: genetic variation and noise-induced phenotypic plasticity. Next, we investigated how the adoption of these two adaptive strategies depends on *c* and *σ* by computing the noise-induced isogenic phenotypic variance, *V*_*ip*_, and the genotypic variance, *V*_*g*_.

### Evolution of robustness and plasticity

We computed *V*_*ip*_ and *V*_*g*_ separately (see *Methods*). The evolutionary time courses of *V*_*ip*_ and *V*_*g*_ are shown in Fig 3. Here, the variances *V*_*ip*_ and *V*_*g*_ corresponding to all output genes and 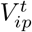 and 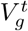 of the target genes are plotted in Fig 3(ii). The variances of the output genes are plotted for different values of the interaction strength *c* and the noise level *σ*. According to the values of *V*_*ip*_ and *V*_*g*_, the four phases are classified as follows.

**Fig 3.**
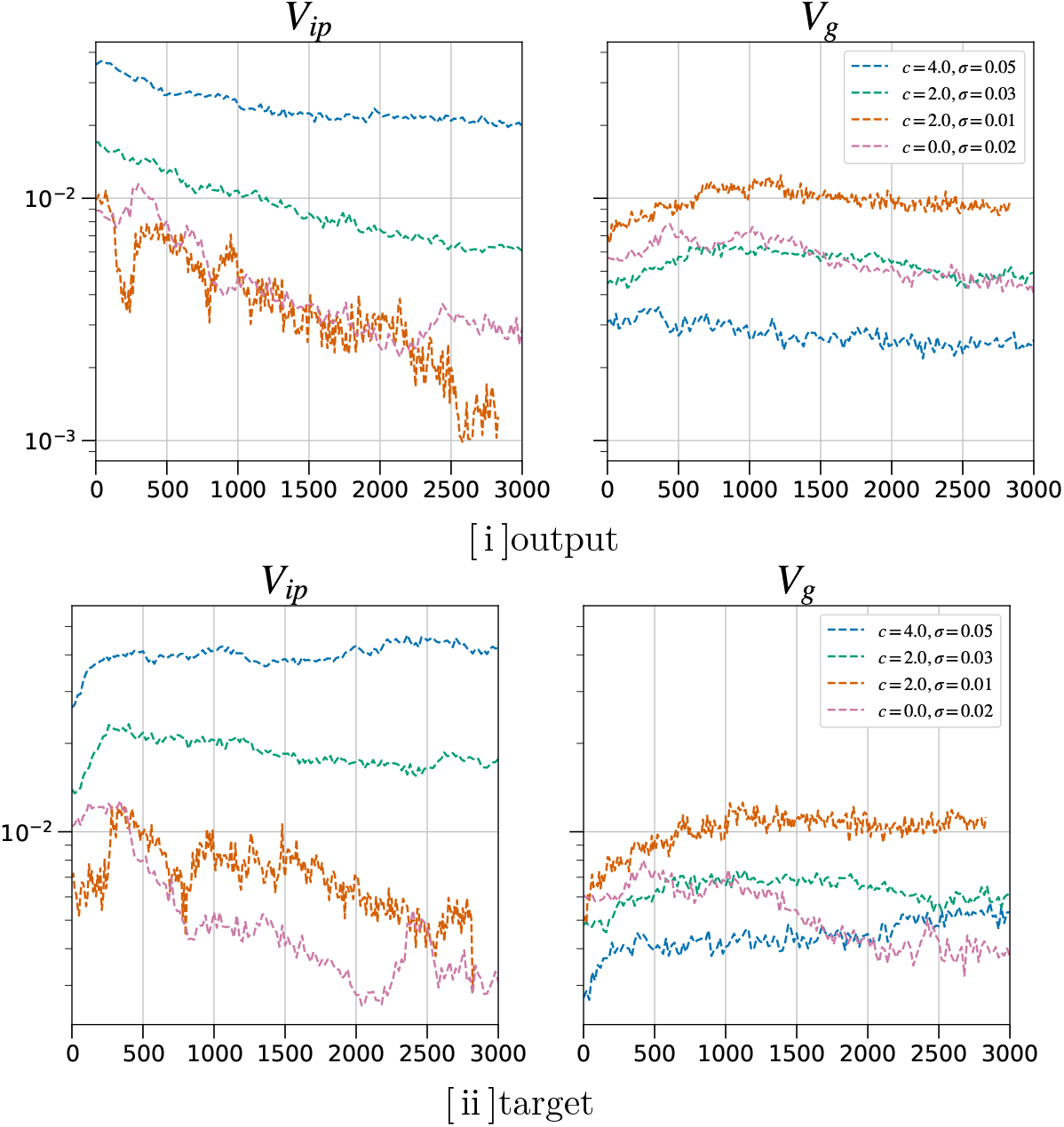
Evolutionary time course of the average phenotypic variances *V*_*ip*_ and *V*_*g*_. The time course of the average phenotypic variances *V*_*ip*_ and *V*_*g*_ (i.e., 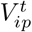 and 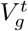). [i] Average *V*_*ip*_ and *V*_*g*_ over all the output genes are plotted against the generation for different values of the noise level *σ* and the interaction strength *c*, as indicated by different colors. Blue: example of evolution with no host-parasite interactions. Hosts are occupied by individuals with the fittest phenotype, thereby losing phenotypic diversity (*σ* = 0.02). Both the isogenic phenotype variance *V*_*ip*_ and the genetic variance *V*_*g*_ decrease. Orange: the case with *c* = 2 and *σ* = 0.01, showing that *V*_*ip*_ decreases, whereas *V*_*g*_ maintains high values, implying the evolution of genotypic diversity. Green: *c* = 2 and *σ* = 0.03. Both *V*_*ip*_ and *V*_*g*_ increase. Red: *c* = 4 and *σ* = 0.05. *V*_*ip*_ decreases slightly, whereas *V*_*g*_ maintains low values. [ii] 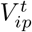 and 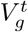 for the target genes. The line colors represent the equivalent conditions as in [i].

#### Phase I: Non-interacting, non-robust (*σ* < *σ*_*c*_ ∼ 0.01 and *c* < *c*_*t*_ = 1)

*V*_*g*_ is maintained at a large value when the noise is below a threshold of *σ*_*c*_ = 0.01 and the interaction is weak (*c* < *c*_*t*_ = 1). Host phenotypes (i.e., gene expression) fluctuate around the fittest type, as shown in Fig 1(i), where most of the output genes are expressed, whereas the variances are sustained at moderate values owing to genetic distribution. A large *V*_*g*_ implies that the phenotypes are not sufficiently robust against mutations.

#### Phase II: Non-interacting, robust (*σ* > *σ*_*c*_ and *c* < *c*_*t*_)

The evolution of robustness against the noise in gene expression dynamics occurs when the noise is above a threshold of *σ*_*c*_ = 0.01 and the interaction is weak (*c* < *c*_*t*_ = 1). In this region, the strength of the interaction is weaker than at the transition point *c*_*t*_ = 1, meaning that the attack by the parasite is not sufficient to cause phenotypic diversification (Fig 2). Figure 4 shows that the values of both *V*_*ip*_ and *V*_*g*_ are low relative to the case with *c* > *c*_*t*_ and *σ* < *σ*_*c*_. As *V*_*g*_ is small, the phenotype is not significantly changed by genetic variation. *V*_*ip*_ and *V*_*g*_ both decrease while satisfying the inequality *V*_*ip*_ > *V*_*g*_, leading to the evolution of robustness to noise and genetic change. This supports previous studies [6, 9, 10]. The host population is both genetically and phenotypically homogeneous, indicating that the output genes are all switched on and exhibit minimal variation with respect to the fittest type.

**Fig 4.**
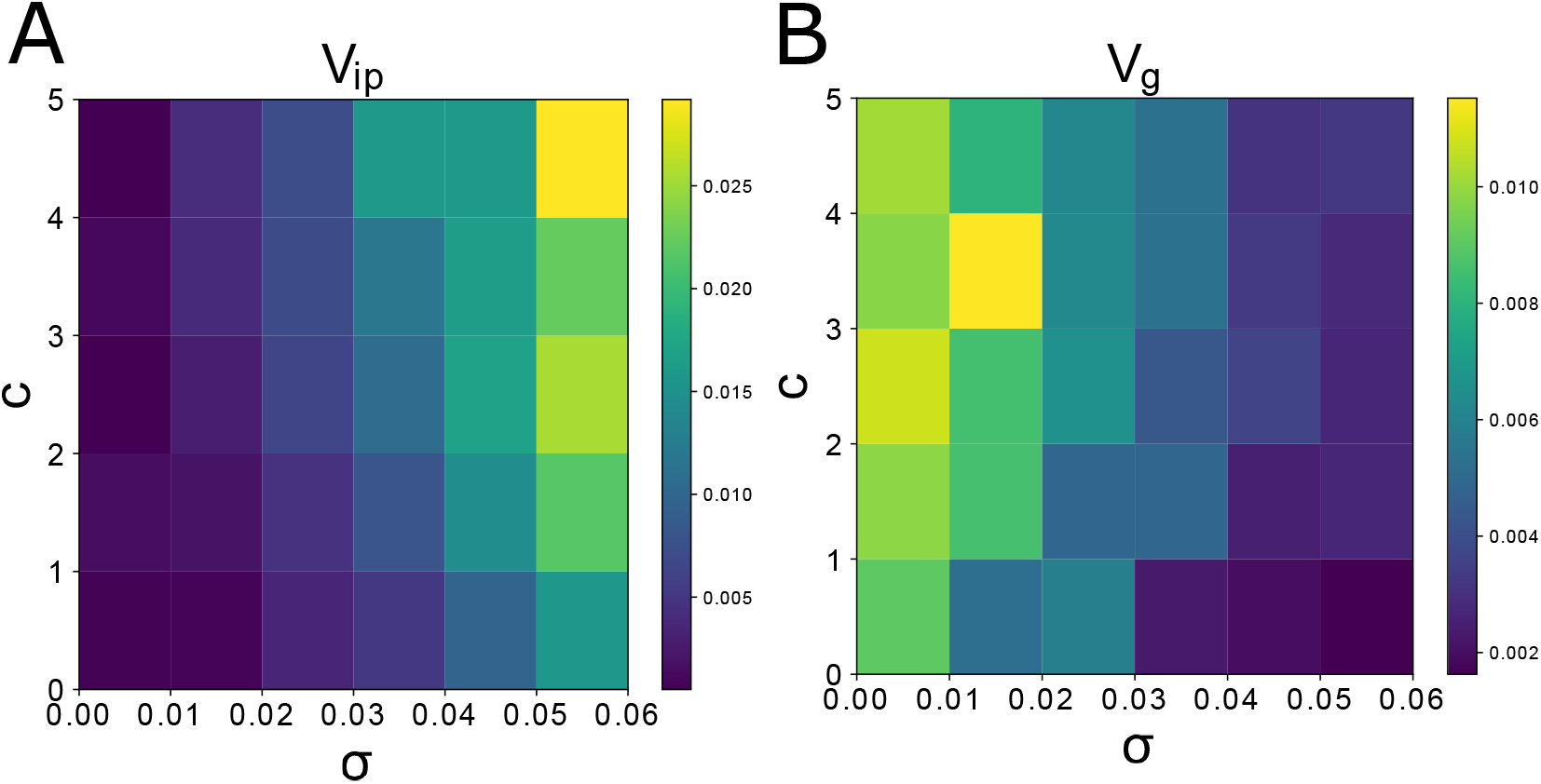
Dependences of *V*_*ip*_ and *V*_*g*_ upon the noise level σ and the interaction strength *c*. *V*_*ip*_ and *V*_*g*_ for all output genes upon the noise level σ (horizontal axis) and the interaction strength *c* (vertical axis). The variance values for each parameter are displayed using color maps. The variance is computed by taking the average of 2500–3000 generations. The color maps indicate that the host-parasite interaction enhances *V*_*ip*_ and *V*_*g*_. Each parameter regime is categorized into one of four phases based on the values of *V*_*ip*_ and *V*_*g*_ (see the text for details).

#### Phase III: Interaction-induced genetic diversification (*σ* < *σ*_*c*_ ∼ 0.01 and *c* > *c*_*t*_)

The evolution of genetic diversification occurs when the noise level is below the threshold *σ* < *σ*_*c*_ and the interaction strength exceeds the transition point *c*_*t*_. No decrease is observed in *V*_*g*_, whereas for *V*_*ip*_, the phenotypic variances remain small, as shown in Fig 5. Phenotypic diversity is generated by genetic diversity. Phenotypes generated from each genotype are concentrated on a unique type such that, for each given genotype, the phenotype does not exhibit plasticity. Multiple groups with different genotypes coexist as a result of interactions with diversified parasites. For each host phenotype, a corresponding parasite exists. The stronger the interaction, the more phenotypes arise, leading to a larger *V*_*g*_ and higher genetic diversity in the population.

**Fig 5.**
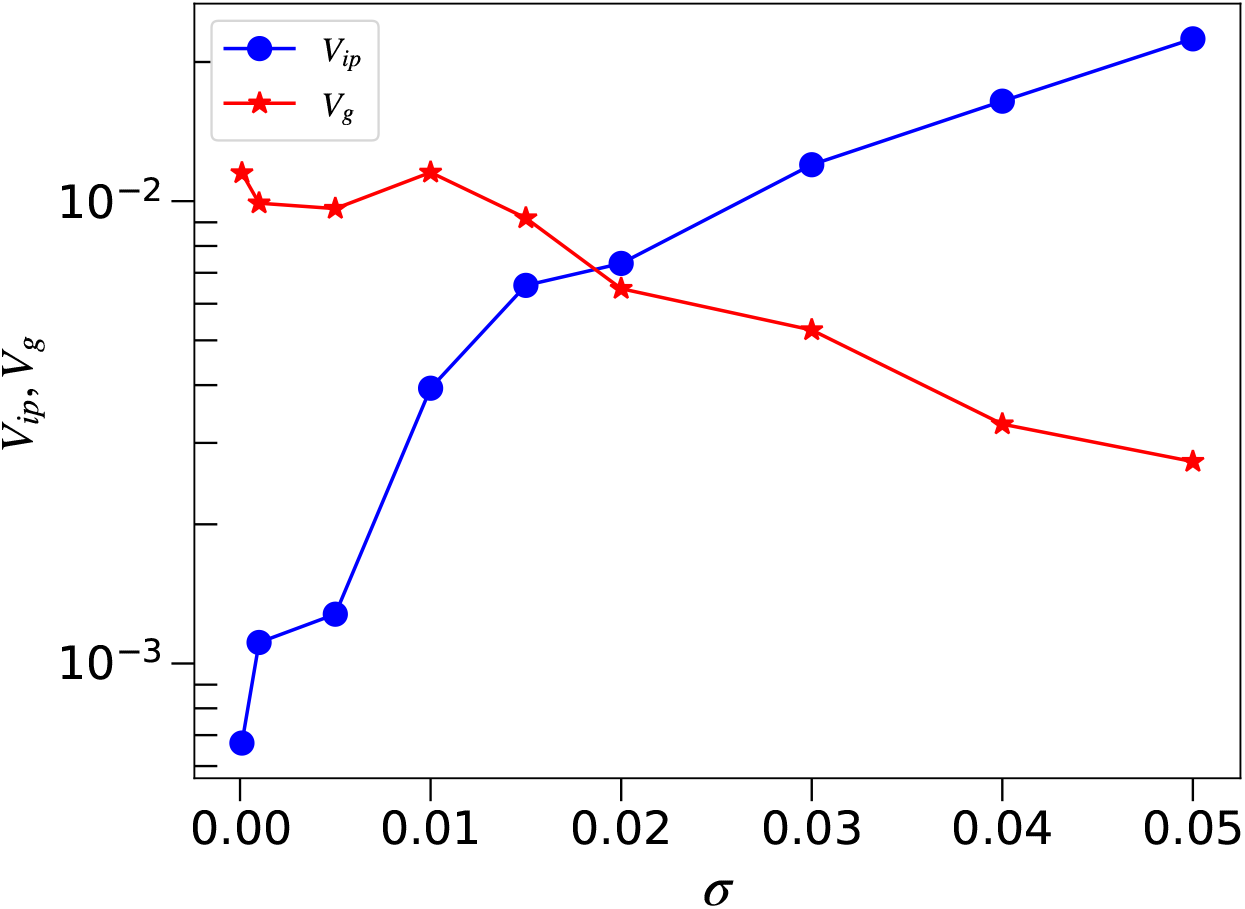
Transition of *V*_*g*_ and *V*_*ip*_ with noise strength. Dependence of the average variances *V*_*ip*_ and *V*_*g*_ of the output genes *i* on the noise level *σ* for a strong host-parasite interaction (*c* = 3). *V*_*ip*_ exceeds *V*_*g*_ at *σ*_*th*_ ≈ 0.01. Based on the data in Fig 4, phenotypic diversification occurs for all parameter values for which the main origin of the phenotypic variances changes from mutation to noise at *σ* ∼ *σ*_*c*_. Beyond the threshold noise level, phenotypic plasticity evolves, with genetic diversification evolving below the threshold.

#### Phase IV: Interaction-induced phenotypic plasticity (*σ* > *σ*_*c*_ and *c* > *c*_*t*_)

For the case of *c* = 3 and *σ* = 0.02, the phenotypic variance of the output gene expression (*V*_*ip*_) increases in the initial stage of evolution (up to 500 generations; see Fig 3). Both *V*_*ip*_ and *V*_*g*_ maintain high values, while satisfying *V*_*ip*_ > *V*_*g*_. Host phenotypes are diversified, without resorting to genetic diversification. Here, isogenic hosts can have more than one phenotype. In addition, gene expression levels are diversified by noise, implying the phenotypic plasticity of the isogenic population. The variances of the parasite-target genes with the parasite are the highest, whereas those of the other output genes are also maintained at high levels (see S1 Fig). Accordingly, the fitness is reduced.

### Transition between the Phase III and IV against noise strength

We also explored the transition between the last two phases. Figure 5 shows the dependence of the variances *V*_*ip*_ and *V*_*g*_ of the output genes on the noise level *σ* for *c* = 3. Below the threshold noise level *σ*_*c*_, *V*_*g*_ exceeds *V*_*ip*_, indicating that the robustness to mutation is diminished. Furthermore, the variability of the host phenotype in response to genetic variation is increased, while the expression levels of the target genes are diversified. When the noise level exceeds *σ*_*c*_, a variety of target gene expression levels are produced. Owing to this fluctuation-induced plasticity in gene expression, hosts can reduce parasite attacks more effectively than when the host phenotype is concentrated on the fittest type.

### Coping with parasites by phenotypic plasticity

In the absence of parasites, the variances *V*_*ip*_ and *V*_*g*_ decrease as the host phenotype adapts to the environment, as described by the fitness condition through evolution (Phase II). Next, we investigated whether introducing parasites after this adaptation enables the evolution of plasticity to progress. Therefore, after the host had adapted to the environment and acquired robustness (i.e., after *V*_*ip*_ and *V*_*g*_ had decreased) we switched the interaction strength from *c* = 0 to *c* = 3. The time course of this evolution is shown in Fig 6. After switching the interaction strength, the host growth rate decreases temporarily before increasing as plasticity evolves (see S4 Fig). The variances *V*_*ip*_ and *V*_*g*_ both increase as in Phase IV. In particular, the phenotypic fluctuations (*V*_*ip*_) in the host population show a notable increase within 100 generations. In addition to the target genes, the fluctuation in the expression level of the output genes also increases. Notwithstanding the evolution of robustness, the evolution of plasticity increases as a result of parasite infection.

**Fig 6.**
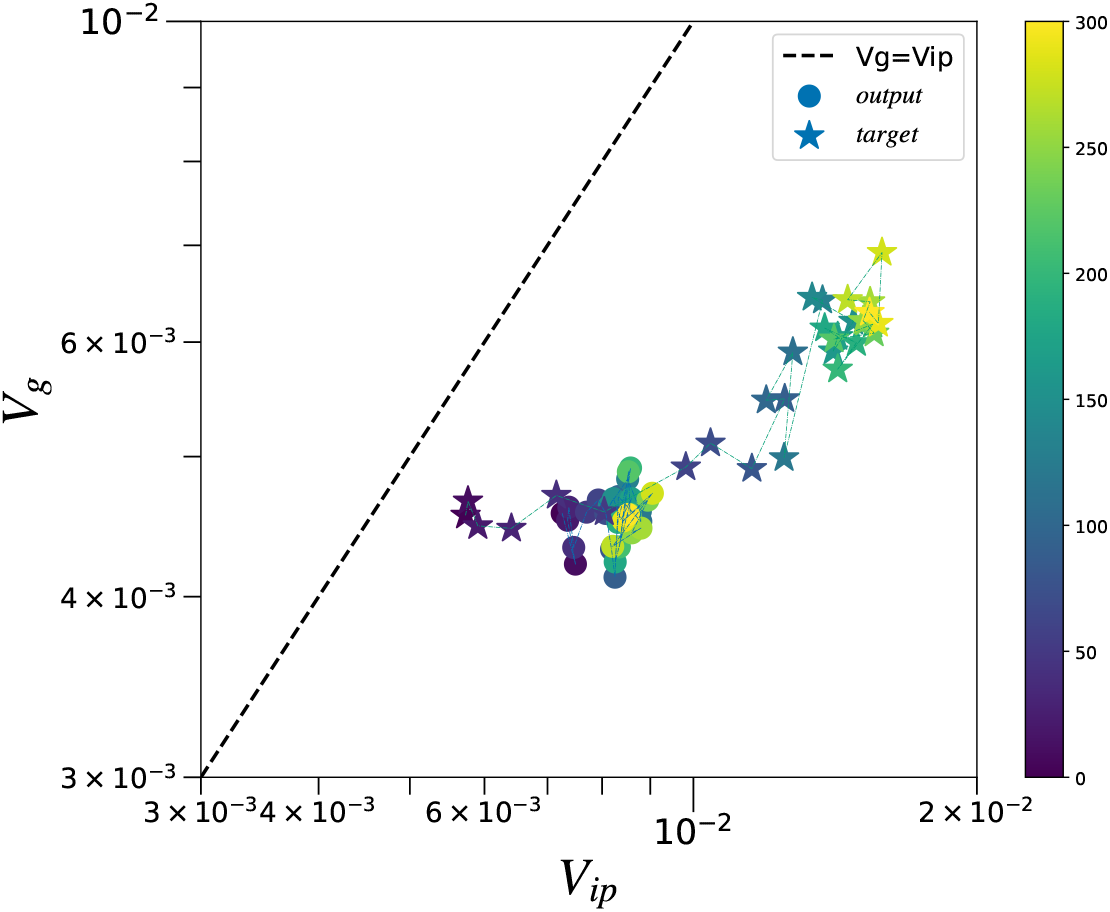
The time course of the variances (*V*_*ip*_,*V*_*g*_) after switching the interaction strength. The time course of the variances (*V*_*ip*_,*V*_*g*_) over generations for *σ* = 0.02. For the first 2000 generations, the evolution of the host was modeled without parasites (*c* = 0). After this, the evolution of robustness was complete (*V*_*ip*_ and *V*_*g*_ decreases). Then, we introduced the interaction with the parasites by changing *c* from 0 to 3. The variances were computed from the isogenic variance over 20 iterations. The time course of 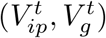 and (*V*_*ip*_, *V*_*g*_) over generations after this switch is plotted for the target (circles; bold lines) and output genes (asterisks; dotted lines). The color bar indicates the number of generations since the switch. The variances 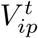 and *V*_*ip*_ both increase owing to interactions with the parasites, resulting in the evolution of phenotypic plasticity. Other model parameters: *N* = 300, *M* = 64, *l*_*inp*_ = 8, *l*_*out*_ = 8, and *l*_*p*_ = 3.

### Dependence of the variances on genetic change and by noise

In a previous study without host-parasite interactions [6], the correlated change between *V*_*ip*_ and *V*_*g*_ during the evolutionary process was observed for *σ* > *σ*_*c*_. Under the pressure of host-parasite interactions, the gene expression dynamics evolved to diversify the phenotypes in the manner described above. Here, we discuss the increase in *V*_*ip*_ and *V*_*g*_ through evolution. As shown in Fig 7(a), the two variance terms pertaining to target gene expression exhibit considerably closer correlation. Moreover, both variances maintain large values, so that the host can escape parasite attacks by producing a different phenotype via mutation. In contrast, Fig 7(b) shows that the variances of other output genes decreased and the evolution of robustness increased for those phenotypes not attacked by parasites. However, this decrease is much smaller than that corresponding to the absence of host-parasite interactions. To increase the phenotypic variance of the target genes, the variances of other gene expressions need to be maintained to a certain degree because of gene interactions through the GRN. This trend of a weak increase in the phenotypic variance for all genes holds for *c* > *c*_*t*_ (see S2 Fig and S3 Fig).

**Fig 7.**
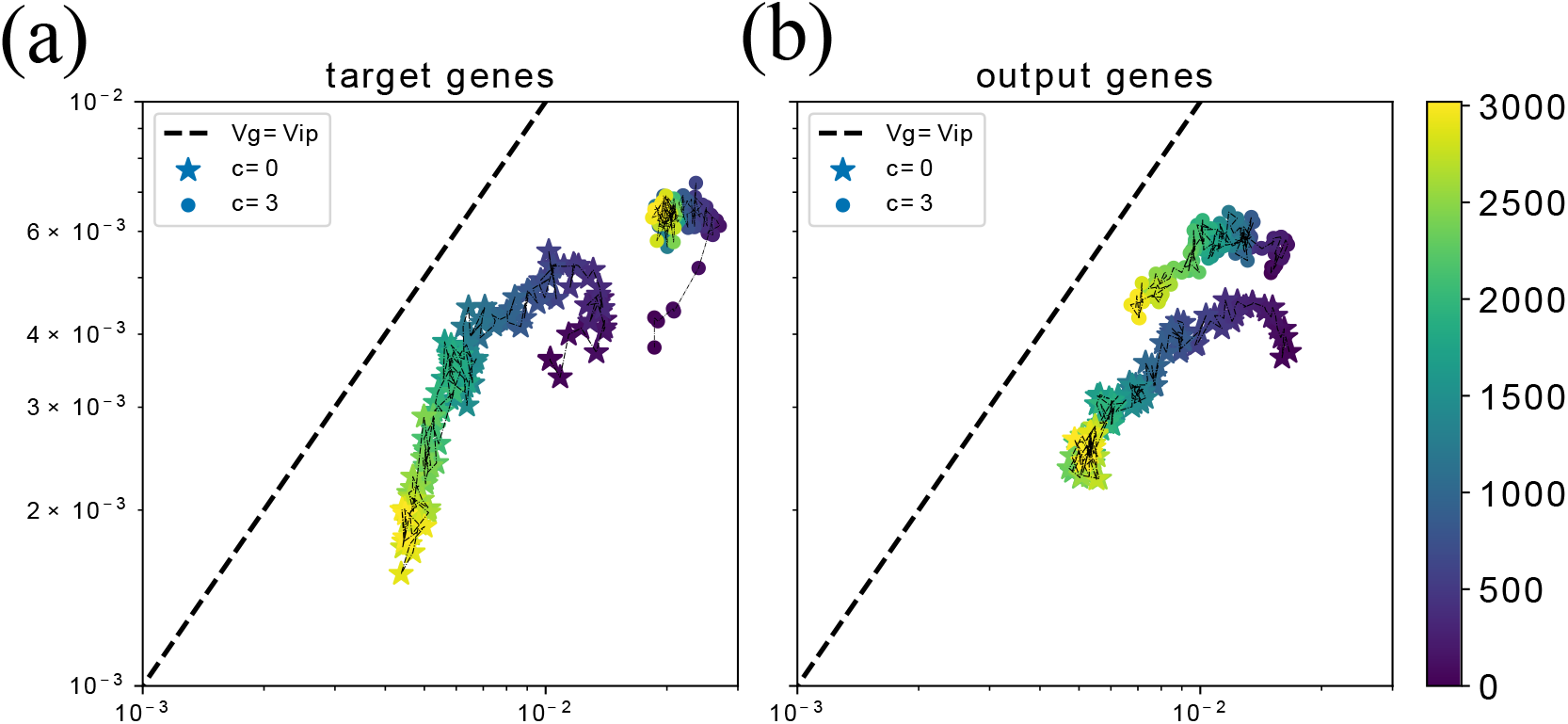
Evolutionary change of the variances for target genes and output genes. The time course of the variance 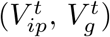 for target genes (a) and the variance (*V*_*ip*_, *V*_*g*_) for output genes (b) (*c* = 0 (asterisks) and 3 (circles)). The variances are computed from the isogenic variance over 100 iterations. The time course over generations is plotted for the variances of gene expression. The plots cover 3000 generations. We set the noise level to *σ* = 0.03 > *σ*_*c*_. In the absence of parasite interactions (*c* = 0), the two variances decrease, while *V*_*g*_ < *V*_*ip*_ is maintained throughout the evolutionary course. Conversely, under host-parasite interactions, *V*_*ip*_ and *V*_*g*_ show a correlated increase. Although this increase is prominent for target genes (asterisks in (a)), it is suppressed for output genes, before showing a slight decrease (b).

### The Trade-off between growth and tolerance

We examined how the increase in the phenotypic variances due to parasite interactions influences the growth rate of a single host. Parasites concentrate their attacks on the hosts with the most frequent phenotype in the population. Figure 8 plots the relationship between *V*_*ip*_ and the fitness, showing the trade-off between the two. Individuals exhibiting high plasticity tend to have lower growth rates owing to the uncertainty in the expression of output genes; however, they are attacked less by the parasites. As the variance *V*_*ip*_ is larger, the parasite attacks are reduced. Consequently, the host types with larger *V*_*ip*_ have a larger chance of survival even if their growth rate as a single cell is lower.

**Fig 8.**
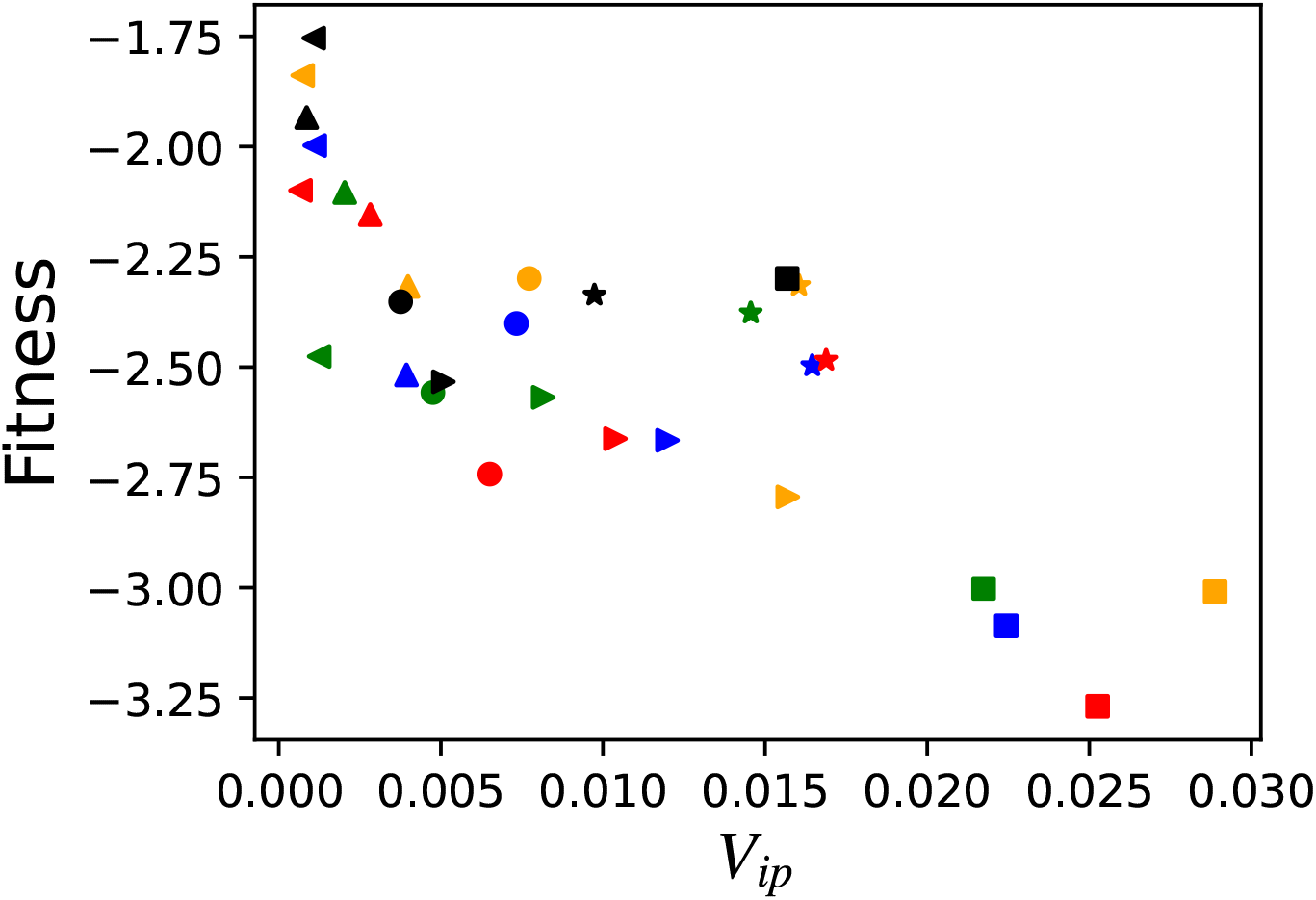
Trade-off between phenotypic plasticity (*V*_*ip*_) and the fitness. Trade-off between phenotypic plasticity (*V*_*ip*_) of the output genes and the fitness 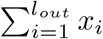 Each values are computed by the average from 2500–3000 generations. Each color represents a different interaction strength: *c* = 0 (black), *c* = 1 (green), *c* = 2 (red), *c* = 3 (blue), and *c* = 4 (yellow). Each marker shape represents a different noise strength: *σ* = 0.05(⋄), *σ* = 0.04(⋆), *σ* = 0.03(▹), *σ* = 0.02(∘), *σ* = 0.01(△), and *σ* = 0.001(◃).

## Discussion

In this study, we studied the evolution of phenotypic variances by using host gene expression dynamics with the regulation network, in the presence of a host-parasite interaction. If the interaction is weak, the host with the fittest phenotype evolves to reduce phenotypic variances. In contrast, if the interaction is sufficiently strong, the phenotypic variances of the host increase to escape specific phenotypes being attacked by each parasite strain. We identified two strategies, either to increase the noise-induced phenotypic variance (*V*_*ip*_) or to increase the genetic variance (*V*_*g*_), depending on the strength of the noise in the stochastic gene expression. If the noise strength is below the noise threshold, the diversification is primarily genetic in origin, whereas above the threshold noise-induced phenotypic plasticity dominates, leading to the variances *V*_*ip*_ and *V*_*g*_ increasing. In the latter case, both variances increase in correlation, thus enhancing phenotypic plasticity, which helps to avoid parasite attacks.

Note that in a fixed environment without inter-species interactions, the two variances *V*_*ip*_ (due to noise) and *V*_*g*_ (due to mutation) tend to decrease, thereby losing phenotypic plasticity and evolvability. Under host-parasite interactions, the GRN evolves to increase these variances, even after robustness has evolved and become established. We classified the diversification of phenotypes into four phases based on the interaction strength *c* and noise level *σ*. The phases were classified according to the respective degrees of phenotypic variance due to phenotypic noise (*V*_*ip*_) and genetic variation (*V*_*g*_).

Whether biological populations deal with environmental changes by genetic variation or phenotypic plasticity has been discussed both theoretically and experimentally [3, 32, 33]. Indeed, the relevance of species-species interactions to phenotypic plasticity and genetic diversification has been discussed extensively [34, 35]. Therefore, it is interesting to examine how the two strategies for phenotypic diversification studied here are adopted therein.

First, we observed a trade-off between phenotypic plasticity and fitness. Hosts that evolved plasticity to deal with parasite interactions tended to exhibit a decrease in inherent fitness (i.e., the growth rate). Interestingly, such a trade-off has been observed for predator-induced phenotypic plasticity [36–38] and in bacteria-phage experiments [39], where the bacteria gain resistance to the phage by reducing the competition for resources. Furthermore, it has also been suggested that excessively strong phenotypic plasticity may reduce adaptability to climate change [40]. In contrast, host populations that exhibit phenotypic diversification by increasing genetic variation (*V*_*g*_) in order to reduce the rate of infection do not show a significant decrease in fitness. However, in this case, their phenotypes are less robust to noise or mutation.

Note that the generation of diverse phenotypes against uncertain environmental changes is known as a” bet-hedging strategy “[41–45]. The question of whether to cope with environmental changes by genetic evolution or by phenotypic plasticity has been discussed in relation to the speed of environmental change. The diversification strategies originating from either *V*_*ip*_ or *V*_*g*_, as discussed here, provide the basis for the increase in phenotypic variance, which is required for” bet-hedging. “

Moreover, the time scale of environmental change is expected to be an important factor in determining which adaptation strategy is chosen. Nevertheless, in the present model, the hosts do not receive parasite information explicitly. The effect of parasite infection on the host is considered only as a negative effect on population growth. The influence of the parasite does not have any effect on the host gene expression dynamics. The characteristics of gene expression dynamics are determined by the noise level *σ*, indicating that the transition point *σ*_*c*_ is independent of the time scale of the parasite population dynamics (see S5 Fig). Furthermore, because the population density is fixed, the interaction strength is constant, meaning that the hosts do not become extinct even if they cannot cope with rapid changes in parasite dynamics. It would be interesting to study the evolution of a model in which the host receives the influence of the parasite directly on the input. For example, in the perceptron model, which produces a favorable phenotype for an input, different strategies emerge depending on the environment [44]. By introducing such parasite inputs, host adaptation strategies are enriched.

In summary, we have demonstrated the evolution of phenotypic plasticity and robustness under host-parasite interactions based on the change in phenotypic variances. Depending on the strength of the interaction and the noise level in the gene expression dynamics, either phenotypic plasticity or genetic diversification evolves. Note that a stronger manifestation of genetic diversification results in speciation. In our forthcoming paper, we aim to show how the phenotypic plasticity induced by host-parasite interactions will also lead to genetic speciation.

## Methods

We define the two phenotypic variances *V*_*ip*_ and *V*_*g*_ as follows.

(1) Gene expression can take different values even among individuals of the same genotype owing to noise during the development process. The variance, denoted as 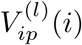, is defined by the variance of 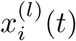 for gene *i* of the *l*-th host-type in the isogenic population, i.e., among these with the same genotype (network 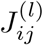).

(2) 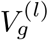 is defined as the phenotypic variance due to the genetic distribution. Here, the gene expression 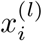 varies both according to the noise and according to the change in the network 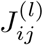. To distinguish the latter from the former, we first computed 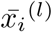, i.e., the average of 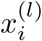 over noise, for each genotype. Then, we computed the variance of 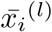 over the heterogenic population by adding a mutation to 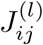. Thus, following the distribution in *J*_*ij*_ (due to mutation), 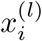 is distributed, with 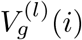 obtained using the variance of 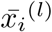 over the heterogenic population. Details on the measurement of each variance are given below. In the simulation, there are up to *N* individuals with different genotypes *J*_*ij*_, with each potentially having a different gene expression. *L* samples are selected at random from the networks. For each selected network, we prepared *N*_*c*_ clones. Then, we computed the variance of the phenotype (gene expression) over the clonal population for each sample at *t* = 1000, after the expression reaches a steady state. The average variance over the sample gives rise to 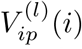, that is, the noise-induced variance over the isogenic population. Then, we prepared 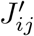 by adding a single mutation to the chosen network *J*_*ij*_ enabling 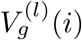 to be estimated for all *L* samples, by applying the same mutation procedure as previously mentioned. We obtained 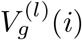 by sampling over a heterogeneous population. We set *L* = 30 and *N*_*c*_ = 50 for all simulations.

## Supporting information

**S1 Fig Average variances of** *V*_*ip*_(*i*) **and** *V*_*g*_(*i*) **(output genes** *i* = 4, 5, …, 8).

**S2 Fig Average variances of** 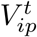 **and** 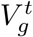 **(target genes** *i* = 1, 2, 3).

**S3 Fig Average variances of** *V*_*ip*_(*i*) **and** *V*_*g*_(*i*) **(other genes** *i* = 9, 10, …, 64).

**S4 Fig Time course of the variances** 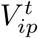 **and the growth rate** *µ* **and** 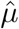.

**S5 Fig Timescale of the parasite changes**.

## Acknowledgments

The authors would like to thank Tetsuhiro Hatakeyama and Nobuto Takeuchi for stimulating discussions on the study. This research was supported in part by a Grant-in-Aid for Scientific Research (A) (no. 20H00123) and a Grant-in-Aid for Scientific Research on Innovative Areas (no. 17H06386) from the Ministry of Education, Culture, Sports, Science and Technology (MEXT) of Japan.

## References

1. de Visser JAG, Hermisson J, Wagner GP, Meyers LA, Bagheri-Chaichian H, Blanchard JL, et al. Perspective: evolution and detection of genetic robustness. Evolution. 2003;57(9):1959–1972.

2. Callahan HS, Pigliucci M, Schlichting CD. Developmental phenotypic plasticity: where ecology and evolution meet molecular biology. Bioessays. 1997;19(6):519–525.

3. West-Eberhard MJ. Developmental plasticity and evolution. Oxford University Press; 2003.

4. Ancel LW, Fontana W. Plasticity, evolvability, and modularity in RNA. Journal of Experimental Zoology. 2000;288(3):242–283.

5. Frank SA. Natural selection. II. Developmental variability and evolutionary rate. Journal of evolutionary biology. 2011;24(11):2310–2320.

6. Kaneko K. Evolution of robustness and plasticity under environmental fluctuation: Formulation in terms of phenotypic variances. Journal of statistical physics. 2012;148(4):687–705.

7. Wagner A. Robustness against mutations in genetic networks of yeast. Nature genetics. 2000;24(4):355–361.

8. Wagner A. Robustness and evolvability in living systems. vol. 24. Princeton university press; 2005.

9. Kaneko K. Evolution of robustness to noise and mutation in gene expression dynamics. PLoS One. 2007;2(5):e434.

10. Kaneko K, Furusawa C. An evolutionary relationship between genetic variation and phenotypic fluctuation. Journal of Theoretical Biology. 2006;240(1):78 – 86.

11. Sakata A, Hukushima K, Kaneko K. Funnel landscape and mutational robustness as a result of evolution under thermal noise. Physical review letters. 2009;102(14):148101.

12. Fisher RA. The genetical theory of natural selection. Oxford University Press; 1930.

13. Futuyma DJ. Evolutionary Biology (2nd ed): by Douglas J. Futuyma. Sinauer Associates; 1987.

14. Hartl DL, Clark AG. Principles of Population Genetics 4th ed. 2007;.

15. Elowitz MB, Levine AJ, Siggia ED, Swain PS. Stochastic gene expression in a single cell. Science. 2002;297(5584):1183–1186.

16. Bar-Even A, Paulsson J, Maheshri N, Carmi M, O’Shea E, Pilpel Y, et al. Noise in protein expression scales with natural protein abundance. Nature genetics. 2006;38(6):636–643.

17. Tsuru S, Ichinose J, Kashiwagi A, Ying BW, Kaneko K, Yomo T. Noisy cell growth rate leads to fluctuating protein concentration in bacteria. Physical biology. 2009;6(3):036015.

18. Kaern M, Elston TC, Blake WJ, Collins JJ. Stochasticity in gene expression: from theories to phenotypes. Nature Reviews Genetics. 2005;6(6):451–464.

19. Furusawa C, Suzuki T, Kashiwagi A, Yomo T, Kaneko K. Ubiquity of log-normal distributions in intra-cellular reaction dynamics. Biophysics. 2005;1:25–31.

20. Mjolsness E, Sharp DH, Reinitz J. A connectionist model of development. Journal of Theoretical Biology. 1991;152(4):429 – 453.

21. Alberch P. From genes to phenotype: dynamical systems and evolvability. Genetica. 1991;84(1):5–11.

22. Pigliucci M. Genotype-phenotype mapping and the end of the ”genes as blueprint” metaphor. Philosophical Transactions of the Royal Society B: Biological Sciences. 2010;365(1540):557–566.

23. Landry CR, Lemos B, Rifkin SA, Dickinson WJ, Hartl DL. Genetic Properties Influencing the Evolvability of Gene Expression. Science. 2007;317(5834):118–121.

24. Hashimoto M, Nozoe T, Nakaoka H, Okura R, Akiyoshi S, Kaneko K, et al. Noise-driven growth rate gain in clonal cellular populations. Proceedings of the National Academy of Sciences. 2016;113(12):3251–3256.

25. Kaneko K. Proportionality between variances in gene expression induced by noise and mutation: consequence of evolutionary robustness. BMC Evolutionary Biology. 2011;11(1):27.

26. Slatkin M, Lande R. Niche Width in a Fluctuating Environment-Density Independent Model. The American Naturalist. 1976;110(971):31–55.

27. Leibler S, Kussell E. Individual histories and selection in heterogeneous populations. Proceedings of the National Academy of Sciences. 2010;107(29):13183–13188.

28. Rivoire O, Leibler S. The Value of Information for Populations in Varying Environments. Journal of Statistical Physics. 2011;142(6):1124–1166.

29. Brockhurst MA, Koskella B. Experimental coevolution of species interactions. Trends in ecology & evolution. 2013;28(6):367–375.

30. Weitz JS, Hartman H, Levin SA. Coevolutionary arms races between bacteria and bacteriophage. Proceedings of the National Academy of Sciences. 2005;102(27):9535–9540.

31. Buckling A, Rainey PB. Antagonistic coevolution between a bacterium and a bacteriophage. Proceedings of the Royal Society of London Series B: Biological Sciences. 2002;269(1494):931–936.

32. Charmantier A, McCleery RH, Cole LR, Perrins C, Kruuk LE, Sheldon BC. Adaptive phenotypic plasticity in response to climate change in a wild bird population. science. 2008;320(5877):800–803.

33. Chevin LM, Hoffmann AA. Evolution of phenotypic plasticity in extreme environments. Philosophical Transactions of the Royal Society B: Biological Sciences. 2017;372(1723):20160138.

34. Agrawal AA. Phenotypic plasticity in the interactions and evolution of species. Science. 2001;294(5541):321–326.

35. Pfennig DW, Wund MA, Snell-Rood EC, Cruickshank T, Schlichting CD, Moczek AP. Phenotypic plasticity’s impacts on diversification and speciation. Trends in ecology & evolution. 2010;25(8):459–467.

36. Lively CM. Predator-induced shell dimorphism in the acorn barnacle Chthamalus anisopoma. Evolution. 1986;40(2):232–242.

37. Imai M, Naraki Y, Tochinai S, Miura T. Elaborate regulations of the predator-induced polyphenism in the water flea Daphnia pulex: kairomone-sensitive periods and life-history tradeoffs. Journal of Experimental Zoology Part A: Ecological Genetics and Physiology. 2009;311(10):788–795.

38. Gordon SP, Hendry AP, Reznick DN. Predator-induced contemporary evolution, phenotypic plasticity, and the evolution of reaction norms in guppies. Copeia. 2017;105(3):514–522.

39. Burmeister AR, Sullivan RM, Lenski RE. Fitness costs and benefits of resistance to phage Lambda in experimentally evolved Escherichia coli. In: Evolution in Action: Past, Present and Future. Springer; 2020. p. 123–143.

40. Oostra V, Saastamoinen M, Zwaan BJ, Wheat CW. Strong phenotypic plasticity limits potential for evolutionary responses to climate change. Nature communications. 2018;9(1):1–11.

41. Simons AM. Modes of response to environmental change and the elusive empirical evidence for bet hedging. Proceedings of the Royal Society B: Biological Sciences. 2011;278(1712):1601–1609.

42. Simons A. Playing smart vs. playing safe: the joint expression of phenotypic plasticity and potential bet hedging across and within thermal environments. Journal of Evolutionary Biology. 2014;27(6):1047–1056.

43. Maxwell CS, Magwene PM. When sensing is gambling: An experimental system reveals how plasticity can generate tunable bet-hedging strategies. Evolution. 2017;71(4):859–871.

44. Xue B, Sartori P, Leibler S. Environment-to-phenotype mapping and adaptation strategies in varying environments. Proceedings of the National Academy of Sciences. 2019;116(28):13847–13855.

45. Beaumont HJ, Gallie J, Kost C, Ferguson GC, Rainey PB. Experimental evolution of bet hedging. Nature. 2009;462(7269):90–93.

